# Water uptake analysis in Japanese mustard spinach: A numerical approach

**DOI:** 10.1101/2019.12.13.875005

**Authors:** Aoi Shimada, Takuya Seki, Musubu Ichikawa

## Abstract

We constructed an experimental system consisting of a syringe pump and water level feedback control to reveal the dynamics of water uptake in Japanese mustard spinach. We also analyzed differences in time-lapse images to quantify turgor movement in these plants. By performing harmonic analysis using the discrete Fourier transform on these dynamics, the frequency domain was obtained. These methods enabled the detailed visualization of plant dynamics.

**Highlight:** In this study, the water uptake dynamics of Japanese mustard spinaches were measured in detail and image difference analysis was performed to investigate the plants’ turgor movements. Frequency domain analysis of the plant dynamics was carried out using a discrete Fourier transform.

## Introduction

In recent years, improving the use of water in agriculture has become essential Therefore, understanding the water dynamics of plants has become important in both agriculture and science. Several studies on the optimization of water use in the field have been reported (Bouman and Tuong, 2001; Tabbal *et al.*, 2002; Smith *et al.*, 2007; Henry *et al.*, 2012; Chen *et al.*, 2018; Takeda *et al.*, 2019*a*, *b*), along with root and plant water uptake models (Steudle and Peterson, 1998; Personne *et al.*, 2003; Wang and Smith, 2004; de Jong van Lier *et al.*, 2013; Lobet *et al.*, 2014; Meunier *et al.*, 2017) In this study, to better understand the water dynamics of Japanese mustard spinach “Komatsuna” (*Brassica rapa var. perviridis*), we constructed an experimental system consisting of a syringe pump and feedback control of the water level. In addition, the turgor movement of the plant was analyzed through image difference analysis of time-lapse images. Harmonic analysis using a discrete Fourier transform (DFT) was calculated from the results and a frequency domain analysis was obtained. The dynamics of water uptake and turgor pressure movement in the plants were visualized in detail.

## Material and method: cultivation of seedlings

Japanese mustard spinach “Komatsuna” (*Brassica rapa var. perviridis*) was cultivated for this experiment. Seeds were sown in a 200mL pot of Sophiterra (KURARAY Co., Ltd.) medium filled with aqueous fertilizer with an EC = 1.5 mS/cm (OAT House No.1, OAT Agrio Co., Ltd.). The seedlings were grown under light emitting diodes (LEDs; (NFSW757 series, Nichia Co.) with a color temperature of 5000 K. The light intensity was adjusted to 150 μmol m^−2^s^−1^of photon flux density. The photon flux density was quantified using a PAR quantum sensor (LI-190R, LI-COR Biosciences). The spectral photon flux density of the LEDs was evaluated with a spectrometer (USB2000+XR, Ocean Optics, Inc.) calibrated with standard calibration light (DH-2000-CAL, Ocean Optics, Inc.) and is shown in Figure 1. The lighting schedule consisted of 12 h light/12 h darkness. The Sophiterra medium was formed of plastic beads with a diameter of ~2 mm and a length of ~3 mm. The seedlings were grown under constant temperature and humidity at a temperature of 21°C and relative humidity of 50%.

**Figure 1.**
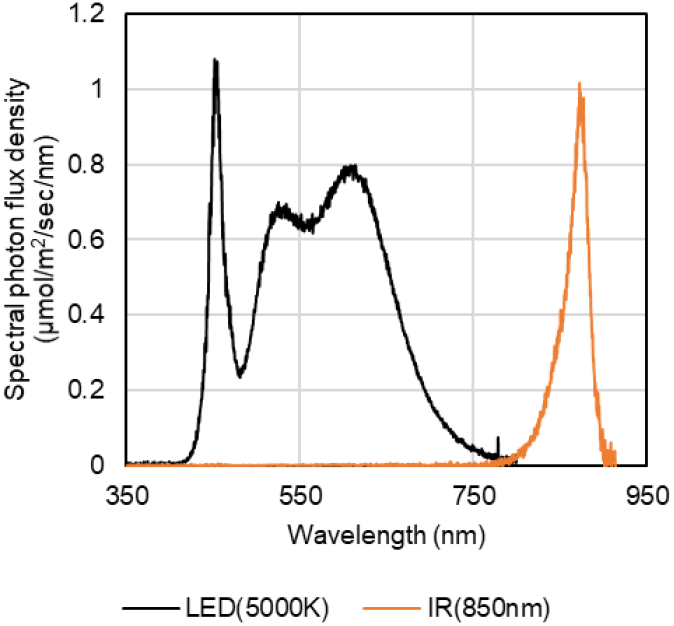
Spectra of LED used in the experiments. A 5000K LED was used as the cultivation light source. Infrared illumination was used for night photography.

### Experiment 1: Water uptake analysis

#### Equipment design and construction

We designed a method to visualize the water uptake in “Komatsuna”. Water in the Sophiterra pot was maintained at constant level using a syringe pump and feedback control system. The water consumption was measured from the control amount in the syringe pump. The syringe pump was designed and built using 3D printing technology (Figures 2 and 3A) for this experiment. The 50 mL syringe (SS-50ESZ, TERUMO Co.) was controlled with a stepping motor (17HS08-1004S, OSM Technology Co.), which was driven by a motor driver IC (DRV8835, Texas Instruments, Inc.). The stepping motor steps were converted to a linear drive using an M3 linear shaft with a 0.5 mm pitch screw. The step angle of the stepping motor was 50 steps/round with a 2-phase induction mode. The syringe gauge was graduated at 1mL/1.5mm. Therefore, the syringe pump could deliver 6.67 μL/step. In addition, the water level reflected the retention of water poured into the Sophiterra medium. A capacitive soil moisture sensor (SEN0193, DFROBOT) was utilized as a water level sensor and inserted into the Sophiterra medium to accurately measure the water level. The water level was converted to a digital value using the 16-bit AD converter (ADS1115, Texas Instruments, Inc.).

**Figure 2.**
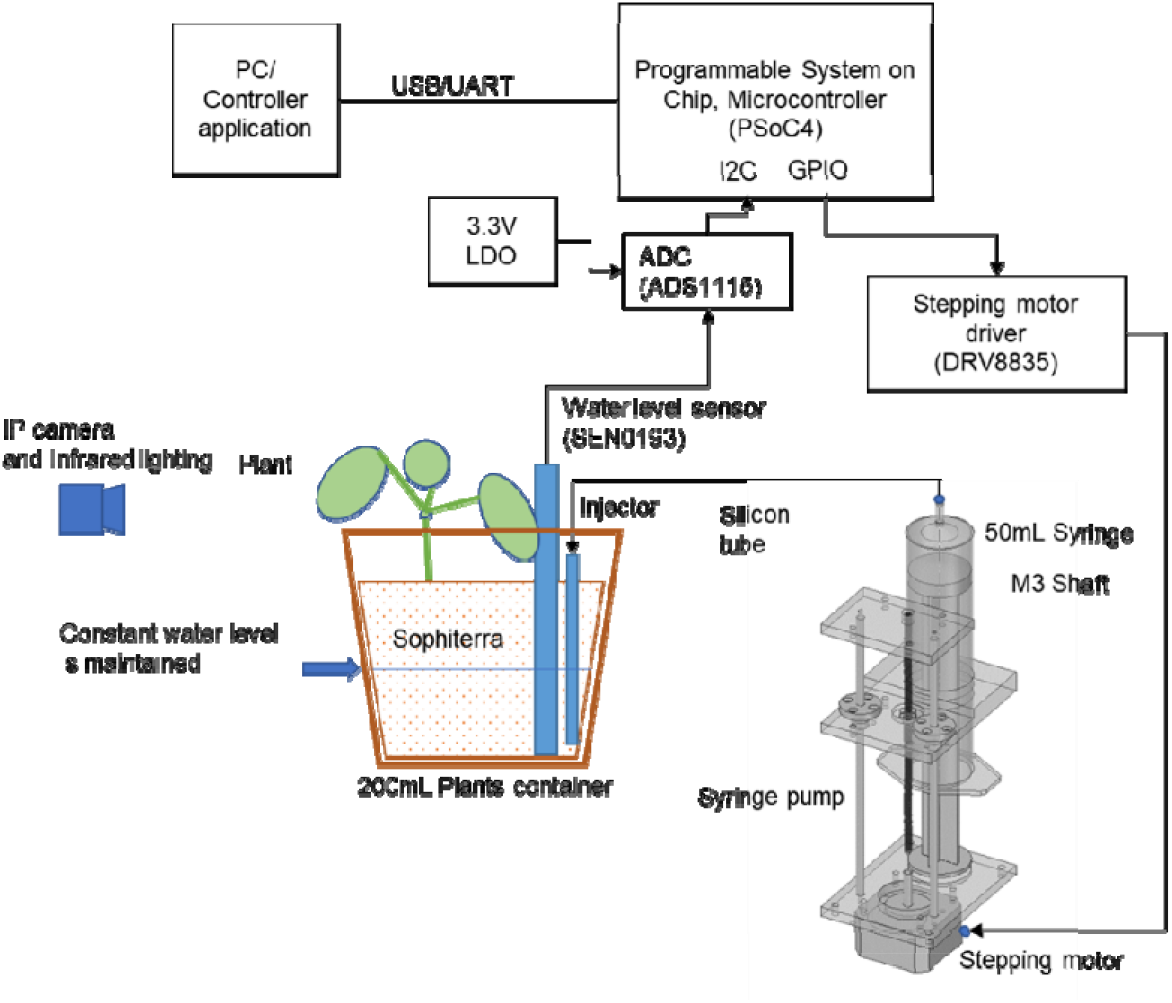
Outline of the experimental system. The plant was equipped with a syringe pump and water level sensor to maintain a constant water 113 level.

**Figure 3.**
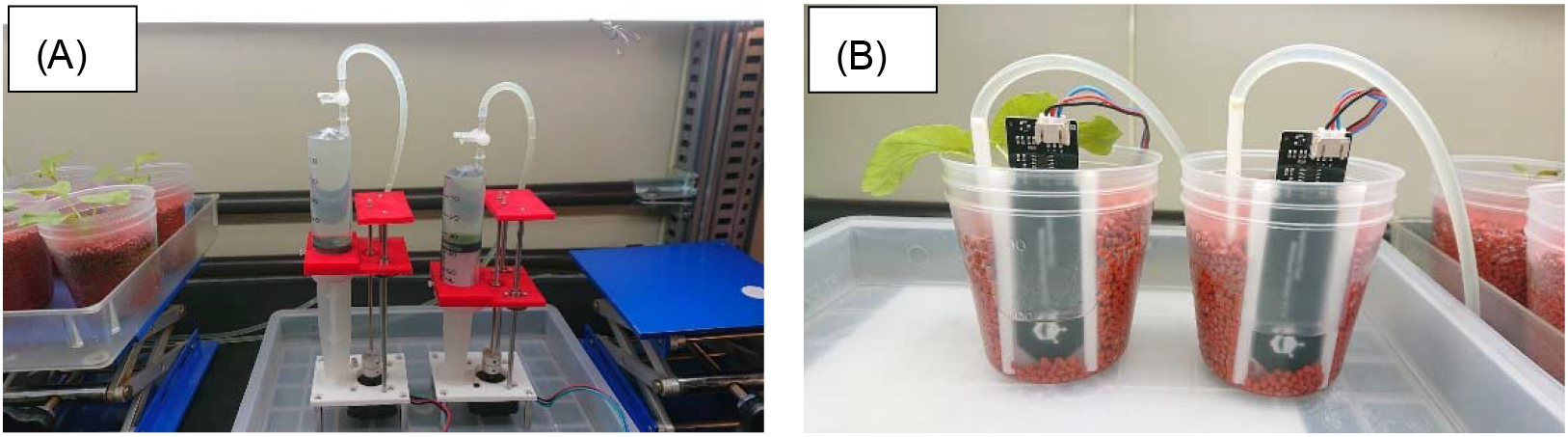
Syringe pump system and a plant. (A) photograph of the syringe pump. (B) a plant equipped with the system.

#### Feedback control system constructed and equipped on a plant

A basic control program was built on a Programable System-on-Chip microcontroller (PSoC4, Cypress Semiconductor Co.). The feedback control system (Figure 2) used to maintain the water level was built on a Windows PC using the C# language.

The water level was measured every 5 s and the feedback control instilled by the syringe pump maintained the water at a constant level. Injector of the water tube was embedded in the bottom part of the pot; thus, water surface was not disturbed by water supply. In fact, the digital value on the analog to digital converter (ADC) connected to the water level sensor was maintained at 24,000 as a specified binary value. The amount of control undertaken by the stepping motor was recorded every 5 min and the cumulative injection of water from the syringe pump was estimated from this record. This system was equipped to a plant on the 23rd day of seedling growth and measurement of water dynamics followed (Figure 3). The water level in Figure 3B was actually obtained by feedback control. A pot containing no plant was prepared at the same time. The plant and system were placed in the constant temperature and humidity room where humidity fluctuation was recorded with a data logger every 5 min. Simultaneously, time-lapse images of the plants were taken.

#### Results and discussion

By measuring the water consumption of the plants and controls precisely, we obtained information about the water dynamics in the plants. Figure 4B, D shows the cumulative injection of water as a function of cultivation days. The water uptake speed was obtained by differentiating (time derivative) the cumulative injection of water and are shown in Figure 4A, B. We applied a moving average of seven frames (boxcar = 7) in three stages to the water uptake speed to remove the noise originating from the system or pump control. Note that the moving average worked as a FIR (finite impulse response) low-pass filter. As the injection speed of water is increasing over approx. 5μL/min, noise from the system was shifted to high frequency and can be subtracted with moving average.

**Figure 4.**
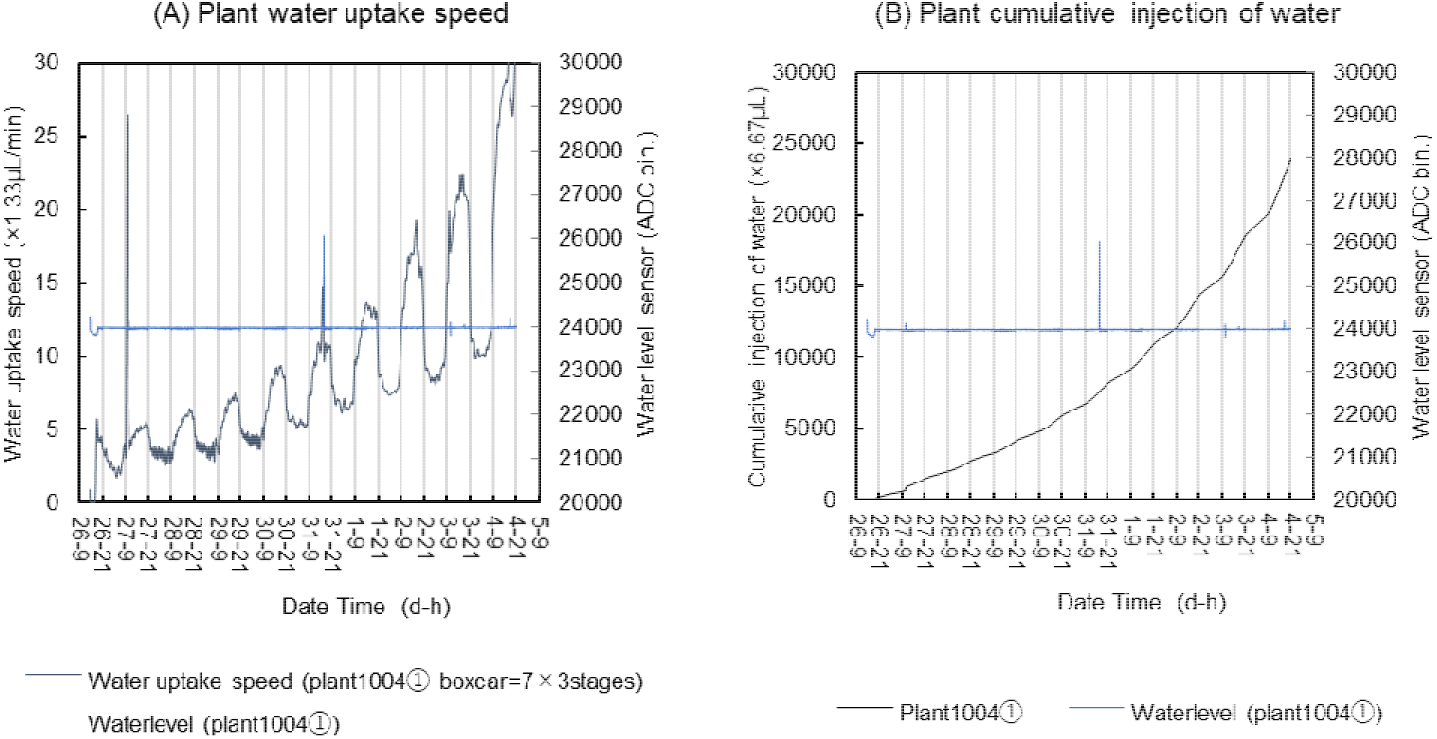
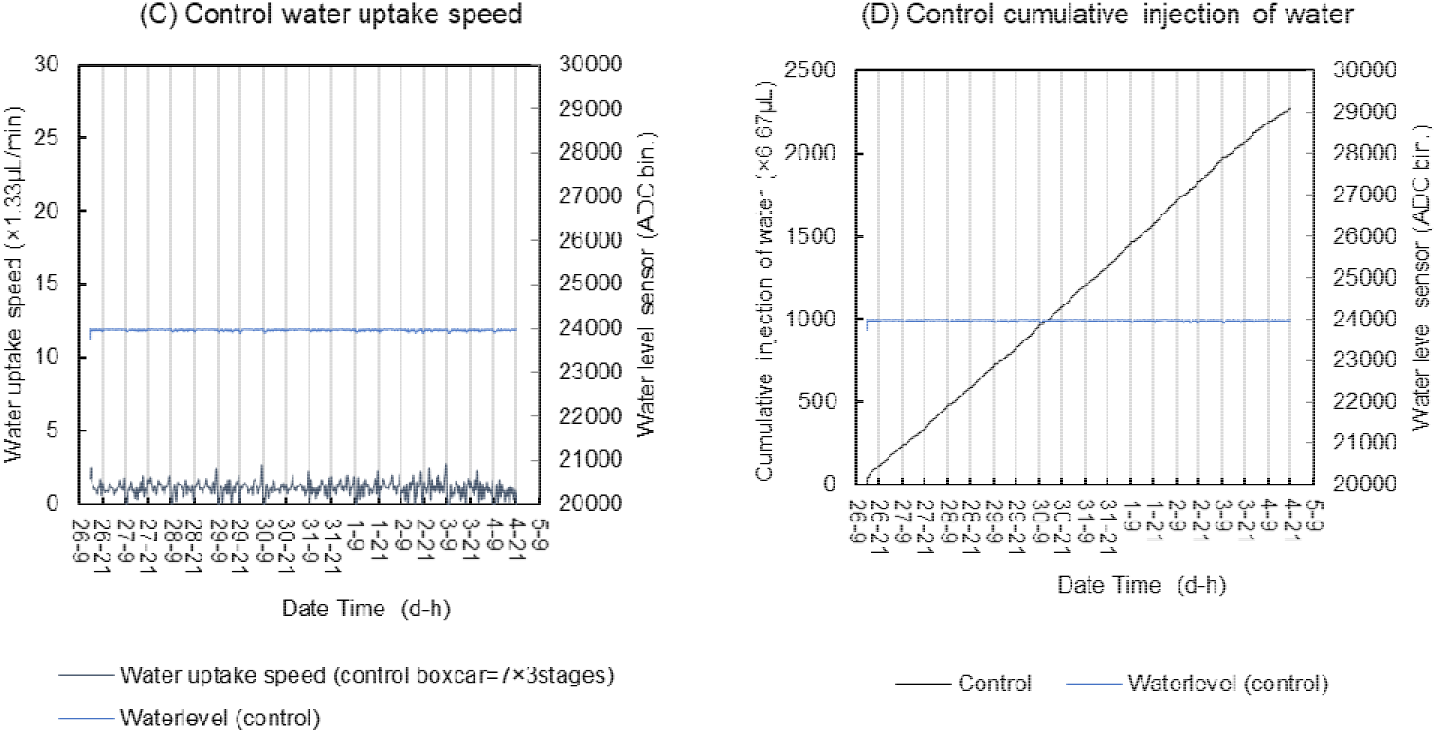
Water uptake analysis of a plant. (A)shows the water uptake speed of the plant estimated from the pump control and smoothed with a moving average (boxcar = 7). A high ripple in the water level corresponded to the timing of the syringe water replacement. (B) shows the cumulative injection of water and (C), (D) show the results for the control.

Figure 4A shows that the water uptake changed greatly between day and night. The water uptake was lower at night and Circadian rhythm (Bünning, 1964) was determined for water flow. The Circadian rhythm is a well-known phenomenon in physiology. (Mohr and Schopfer, 1992; Taiz and Zeiger, 2002). Transience in water uptake may relate to an increase in stomatal conductance during the daytime and suggests that atmospheric humidity may affect water uptake speed. In addition, as the plant grew, the water uptake increased. However, water uptake at night was lower than that at day during all stages of growth. (Figure 4A) Understanding the water flow dynamics of these plants is important to improving agricultural water use and future detailed analyses of these results may offer useful information to breeding programs for creating plant varieties with improved water use.

### Experiment 2: Image difference analysis

Image analysis of plants has been actively undertaken in a number of studies (Nilsson, 1995; Jiazhi and Yong, 2008; Gupta *et al.*, 2014) and, recently, time domain imaging has been studied. (Fukuda *et al.*, 2013; Yoshida *et al.*, 2018; Fukuda, 2019) In this study, in order to quantify and analyze the movement of the whole plant during a time domain, we performed differential analysis of time-lapse images. Time-lapse images were taken every 5 min, considering sampling theorem. A Network IP camera (BB-HCM511, Panasonic Co.) was used in the experiments. Infrared illumination at a wavelength of 850 nm (VSMY98545, Vishay Intertechnology, Inc.) was used to take photographs of the plant continuously day and night. (Figure 1) Infrared light was lit only during the night. The difference of the images was analyzed by taking the difference in each image and obtaining the summation of the total pixels in the difference image Equation 1: where Image(n) is a pixel map of the image, n is number of orders of shooting the image, and Diff(n) is image difference obtained.

Difference analysis is a simplified image analysis method but it can be expected that the movement of the whole plant, including minute and fine movements can be quantified using this technique. The difference analysis of the images was constructed using the OpenCV (Bradski, 2000) open-source image processing engine. In this experiment, OpenCVsharp3 (shimat, 2015) was used as a wrapper build of OpenCV for the C# language platform. First, all the images were converted to grayscale and the difference analysis was calculated for each 5 min period. For the results, a moving average of six frames (boxcar = 6) was utilized. The results of the image analysis are shown in Figure 5.

**Figure 5.**
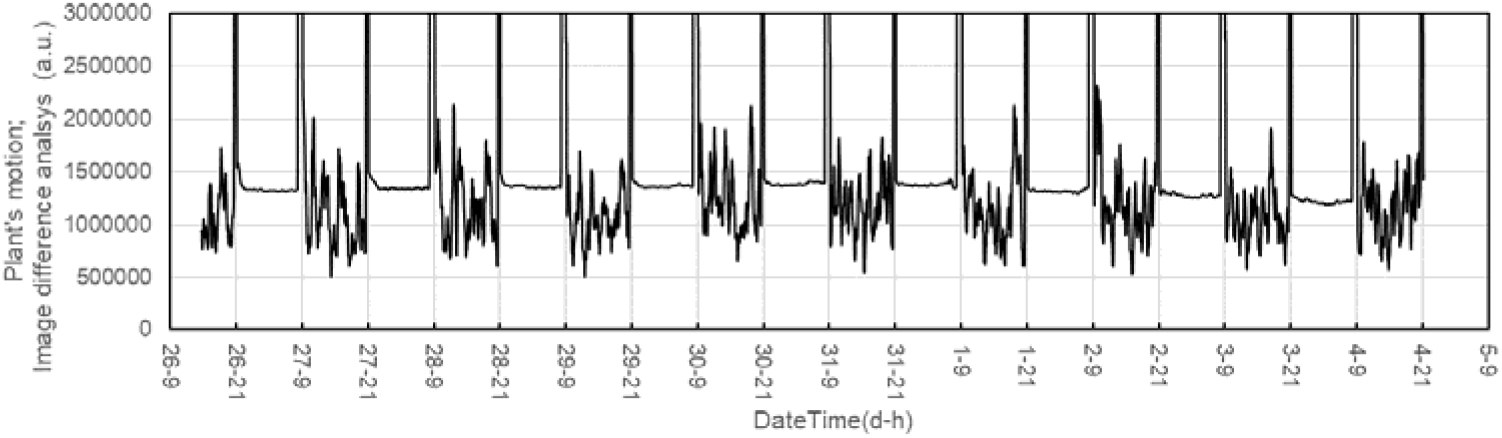
Image difference analysis calculated according to the plant’s motion. Image difference analysis was calculated using OpenCV and the results were smoothed with a moving average (boxcar = 6).

### Experiment 3: Harmonic analysis

Harmonic analysis using a discrete time Fourier transform (DFT) (Press *et al.*, 1992) was carried out for water uptake speed data point sets for every single day and night and the results from the quantification of plant turgor movement. The fluctuation of humidity in the room was also evaluated using a DFT.

The DFT was performed on the results from each day and night for 5 days from the 27th day after sowing and the average of the results was obtained. Thereby, the characteristics of the frequency distribution were evaluated. The Fourier transform was calculated by the DFT with the Blackman Harris window function. (Harris, 1978)(MathWorks, 2019). The Blackman Harris window has small side lobes and is suitable for detecting minute signals. For plant turgor movement analysis and water uptake, DFT was calculated after the moving average (boxcar = 6 or 7) was applied to the time domain data.

#### Results and discussion

As the result of the Fourier transform, the frequency domain analyses were obtained. These analytical series are shown in Figure 6. It was found that the water uptake included a fluctuation at ~0.8 h. The fluctuation of the relative humidity in the constant temperature and humidity chamber also showed a fluctuation at ~0.8 h and it can be seen that the uptake of water was correlated with relative humidity. It is suggested that stomatal conductance is related to the water uptake.

**Figure 6.**
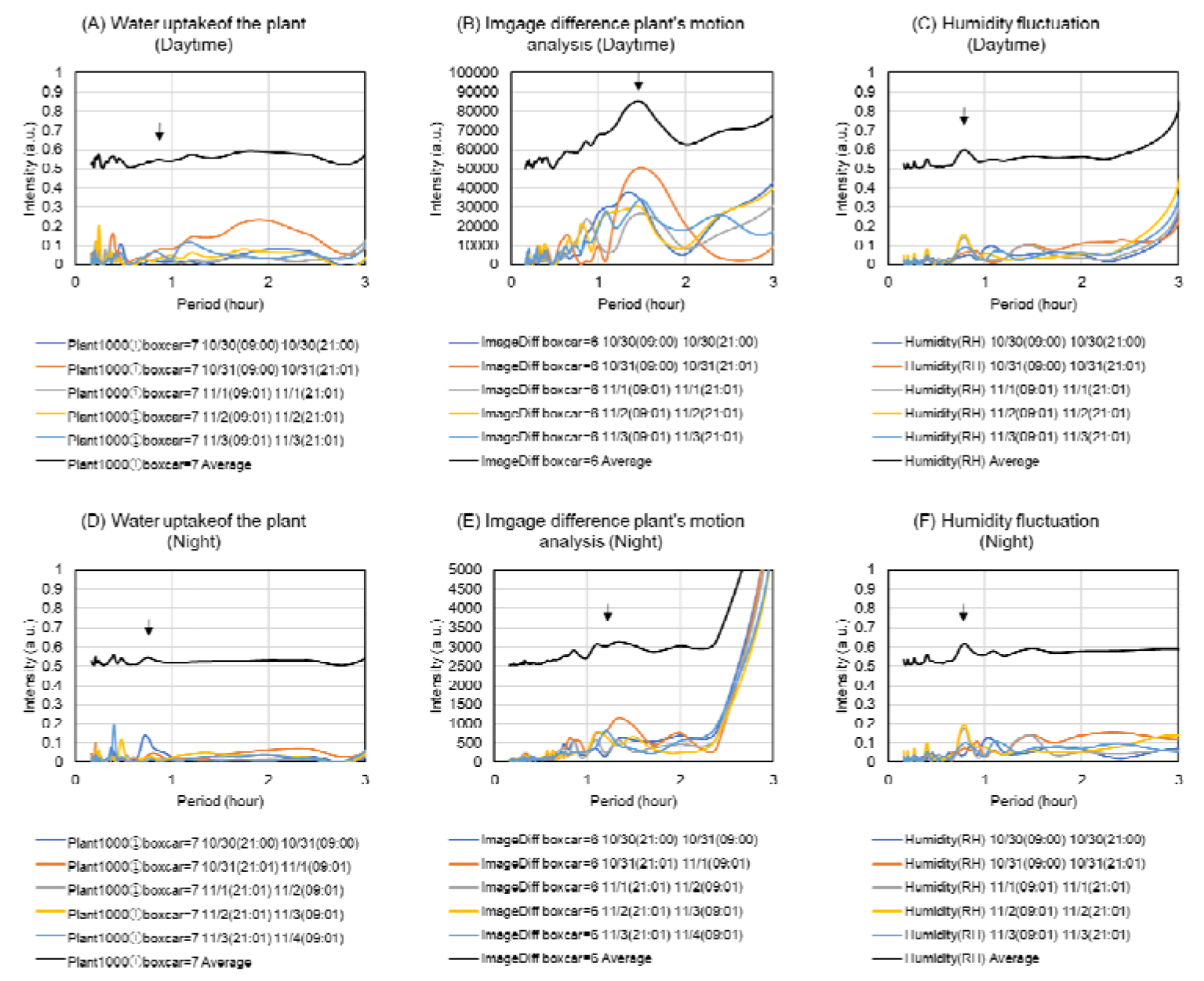
Harmonic analysis using the DFT. Harmonic analysis was calculated with the DFT. (A, D) shows the DFT analysis for water uptake speed, (B, E) is the DFT for the image difference analysis and (C, F) is the fluctuation of relative humidity in the room. (A), (B), and (C) all show the results for the daytime. However, (D), (E), and (F) show the results for the night.

However, the image difference analysis, including the plant motion data, showed a characteristic frequency component of ~1.5-h (Figure 6B, E). Here, it should be noted that the difference analysis of the image was evaluated as an absolute value. Thus, plant movement was evaluated as a positive value regardless of whether it was up, down, left, or right. This, therefore, contained a frequency that was twice the function of the plant leaf position. Therefore, from the 1.5-h period in the image difference analysis, it was estimated the fluctuation at the 3-h period was a function of plant leaf position. There was no significant correlation between water uptake and turgor movement. It should also be noted that the image difference analysis data include not only the plant’s vibrations but also the frequency of the change in growth rate. Moreover, when comparing each of the obtained harmonic analysis series, the characteristics of the frequency may differ depending on whether it was day or night. It can be assumed that this is a result of superimposing pressure adjustment and various regulations depending on the growth stage.

## Summary

In this study, the dynamics of water uptake of the Japanese mustard spinach were revealed in detail by constructing an original experimental system. Moreover, turgor movement of the plant was quantified by image difference analysis. These analyses unveiled the dynamics of the plant in detail. We then used a numerical approach involving harmonic analysis using a DFT to obtain a frequency domain analysis. We determined motion related to the fluctuation in relative humidity and other periodic phenomena. The relationship between stomatal conductance and growth stage and other regulatory events in the plants was suggested.

In future studies, we will investigate the effects of light and related plant physiology. We believe that this work will assist in the improvement of water use efficiency in plants. Moreover, understanding these plant physiological dynamics will contribute to the development of various biosensor applications.

## Acknowledgement

First, we would acknowledge for Prof. Hatsumi Nozue of Shinshu University in advice on plant physiology.

We acknowledge for Ms. Chiaki Okamoto, Ms. Yayoi Karaki and Dr. Kana Shirai of Shinshu University, SU-PLAF, Research Center for Advanced Plant Factory about using of the constant temperature and humidity room and center’s facilities.

We would acknowledge for Prof. Nobuaki Hayashida of Shinshu University in discussions on cruciferous plants. And We would also acknowledge Prof. Masayuki Nozue and Prof. Masahiro Nogawa of Shinshu University for discussions on water use in agriculture and plant physiology.

